# Dynamic Cell Imaging: application to the diatom *Phaeodactylum tricornutum* under environmental stresses

**DOI:** 10.1101/2021.10.22.465453

**Authors:** Houda Bey, Florent Charton, Helena Cruz de Carvalho, Shun Liu, Richard G. Dorrell, Chris Bowler, Claude Boccara, Martine Boccara

**Affiliations:** Institut Langevin, ESPCI Paris, PSL Research University, CNRS UMR 7587, 1 rue Jussieu, 75005 Paris, France; Ecole Normale Supérieure, PSL Research University, Institut de Biologie de l’Ecole Normale Supérieure (IBENS), CNRS UMR 8197, INSERM U1024, 46 rue d’Ulm, F-75005 Paris, France; Université Paris Est-Créteil (UPEC), Faculté des Sciences et Technologie, 61, avenue du Général De Gaulle 94000 Créteil, France

**Keywords:** Chloroplast, Endoplasmic reticulum, Light scattering, Lipid droplets, Iron limitation, Organelle movement, Phosphate limitation

## Abstract

The dynamic movement of cell organelles is an important and poorly understood component of cellular organisation and metabolism. In this work we present a non-invasive non-destructive method (Dynamic Cell Imaging, DCI) based on light scattering and interferometry to monitor dynamic events within photosynthetic cells using the diatom *Phaeodactylum tricornutum* as a model system. For this monitoring we acquire few seconds movies of the signals that are related to the motion of dynamic structures within the cell (denoted scatterers), followed by a statistical analysis of each pixel time series. Illuminating *P*.*tricornutum* with LEDs of different wavelengths associated to short pulsed or continuous-wave modes of illumination revealed that dynamic movements depend on chloroplast activity, in agreement with the reduction in the number of pixels with dynamic behaviour after addition of photosystemII inhibitors. We studied *P. tricornutum* under two environmentally relevant stresses, iron and phosphate deficiency. The major dynamic sites were located within lipid droplets and chloroplast envelope membranes. By comparing standard deviation and cumulative sum analysis of the time series, we showed that within the droplets two types of scatterer movement could be observed: random motions (Brownian type) but also anomalous movements corresponding to a drift which may relate to molecular fluxes within a cell. The method appears valuable for studying the effects of various environments on a large variety of microalgae in the laboratory as well as in natural aquatic environments.

**HIGHLIGHTs:** Light scattering an alternative to fluorescence to rapidly evidence dynamic processes.

Lipid droplets the major metabolic active sites under stress

A non-destructive visualisation method for laboratory microalgae and aquatic samples..

**SIGNIFICANCE STATEMENT:** Light scattering could be an alternative to fluorescence techniques to study dynamic processes within photosynthetic cells. We used a method combining light scattering and interferometry to analyse movements of intracellular scatterers in the marine diatom *Phaedactylum tricornutum* under two environmentally relevant stresses, iron and phosphate deficiency. Lipid droplets were the major active sites under stress. The method which is rapid and non destructive can be broadly expanded to study other microalgae and their stress responses, in the laboratory and in aquatic environments.

## INTRODUCTION

Diatoms are a diverse group of photosynthetic microorganisms, which account for up to 40% of ocean primary production (Bowler *et al*., 2010). They are distributed worldwide, from tropical and subtropical regions to polar ecosystems in oceans and fresh waters, and thus have an exceptional ability to adapt to highly dynamic aquatic environments (Falciatore *et al*., 2020; Pierella *et al*., 2020). These include stressful environmental fluctuations and chronic scarcities of key nutrients including nitrogen, phosphorus or iron (Abida *et al*., 2015 ; Alipanah et al., 2015; Cruz de Carvalho *et al*., 2016; Kazamia *et al*., 2018; Gao *et al*., 2021), extreme temperatures ranging from continuous near-freezing temperatures in ocean polar regions to hot currents in equatorial oceans (Yao *et al*. 2015; Liang *et al* 2019) or high light illumination levels (Domingues *et al*., 2012; Alboresi *et al*., 2016).

Diatoms have a complex evolutionary origin, which involved a secondary symbiotic event where a heterotrophic eukaryote engulfed a microalga of the red lineage (Cavalier-Smith, 1999). This has led to the presence of four membranes around the plastids of diatoms, with the outermost chloroplast membrane joined to the endoplasmic reticulum (ER) and several internal chloroplast compartments including the periplastid compartment (located between the second and third innermost membranes). To identify the metabolically active organelles in diatom cells under stress, methods involving organelle isolation, biochemical methods, genomic, and cell products characterisation have been very valuable (Abida *et al*., 2015; Lupette *et al*., 2019). Fluorescence imaging techniques with high spatial and temporal resolution have been previously used to further investigate dynamic processes in stressed diatoms (Lupette *et al*., 2019; Jaussaud *et al*., 2020). Indeed, cytological methods involving the use of specific stains or of fluorescent proteins are widespread methods used in microscopy. Green Fluorescent Protein (GFP) tagged proteins are used to visualize and track in real time proteins of interest as well as cell organelles (Apt *et al*., 2002; Kazamia *et al*., 2018). Among the many GFP applications, Fluorescence Recovery After Photobleaching (FRAP) has been used to characterize the transport of integral membrane proteins from the endoplasmic reticulum to lipid droplets in yeasts (Jacquier *et al*., 2011).

Studies of biological materials by Dynamic Cell Imaging (DCI) also called Dynamic Full field Optical Transmission Tomography (D-FFOTT) are based on the detection of light back-scattered generated by subcellular organelles (denoted scatterers) movements and do not need elaborate molecular manipulations, as is the case for GFP. This method is non-invasive and non-destructive (Apelian *et al*., 2016, Thouvenin *et al*., 2017, Scholler *et al*., 2019). A more recent variation of DCI, working in transmission and forward scattering,that we use here is described in Thouvenin *et al*., 2021. DCI was shown to be associated with metabolic activity as it disappeared when cells were fixed or treated with the metabolic inhibitor deoxy-glucose (Apelian *et al*., 2016). DCI appeared thus as a promising technique to study cell substructure dynamic in eukaryotic microorganisms unable to be genetically modified. For that reason we first studied the model unicellular photosynthetic microorganism *Phaeodactylum tricornutum* responding to nutrient stress conditions.

We studied two major nutrient stresses: iron and phosphate depletion. Iron (Fe) is an essential micronutrient for of all living cells since it is crucial for photosynthesis and respiration, and for phytoplankton it is a cofactor of proteins involved in a number of metabolic reactions. Iron deficiency leads to a decrease in photosynthetic efficiency (Kolber et al., 1994) by inducing partial blocking of the electron transfer between photosystem II (PSII) and photosystem I (PSI) (Roncel *et al*., 2016). Iron deficiency is an important growth-limiting or co-limiting nutrient in many regions of the world’s ocean in which diatoms are important primary producers (Ustick *et al*., 2021).

Phosphate is a fundamental element of all living cells being part of membranes, nucleic acids and other biomolecules. Phosphate limitation and co-limitation is frequently observed in low northern latitudes (Ustick *et al*., 2021). The deficiency of its inorganic form (Pi) is also a recurrent form of stress that highly limits ocean primary production. Indeed, inorganic phosphate (Pi) deficiency is known to induce a shift in diatom metabolism leading to arrested cell growth and production of lipid vesicles (Cruz de Carvalho *et al*., 2016). It has recently been shown that these lipid vesicles (also known as lipid bodies or lipid droplets, accumulate in the vicinity of the chloroplasts (Alipanah *et al*., 2018; Lupette *et al*., 2019). Indeed limitation of nutrients such as nitrate and phosphate leads to remodelling of cellular membranes, with degradation of membrane phospholipids and fatty acids which are redirected to lipid bodies (Goss *et al*., 2020).

Various possible kinds of organelles movements can be observed in cells from a pure Brownian type to drifts with a defined velocity (Saunter *et al*., 2009). Here we characterized the slight amplitude movements of scatterers within *P. tricornutum* by analysing the time series associated to each pixel of the field of view in order to get a map of the transmission, the standard deviation, the cumulative sum and the average frequency. Through this computational analysis, we were able to highlight random and non-random driven movements suggesting that this method can spot *in organello* molecular fluxes.

## Material and Methods

### Diatom culture conditions

*Phaeodactylum tricornut*um CCMP632 was cultured in artificial sea water (40 g/L, Sigma) supplemented with f/2 nutrients and vitamins (F/2 Media Kit Bigelow NCMA) under continuous shaking (100 rpm) at 20 °C under cool white fluorescent lights (30 µE.m^-2^ s^−1^) with a 12 h photoperiod. For phosphate depletion studies equal aliquots of 4 day-old cultures from the same batch culture were inoculated in parallel in fresh f/2 media (control conditions) and in 250 mL fresh f/2 media without phosphate supplement (Pi depleted) and cultured in the same conditions as described above for 8 days.

For iron depletion, cultures were grown in enhanced seawater medium with or without iron-EDTA (supplemented concentration 86.9 µmol/ l), and analysed after two subcultures (with starting inoculum of 100,000 cells/ml for 8 days. All cell lines were grown in ventilator-capped plastic culture flasks (Sigma) to minimise residual iron or phosphorus contamination from the culture flask into the growth media. Cells were collected by centrifugation and mixed with melted 1% agar in culture medium and immediately observed after jellification.

For photosystem II inhibition, we used 3-(3,4-dichlorophenyl)-1,1-dimethylurea (DCMU; 40 µM) and hydroxylamine (HA; 2mM) solutions in water. Cells were incubated for 10 minutes at room temperature with inhibitors, embedded in agar and immediately observed.

### DCI: Data acquisition and treatment

To follow the movement of internal structures within diatoms we used at the pixel level, the time variation of the optical tomographic images, the principle of which is described in detail in Apelian *et al*., (2016), Thouvenin *et al*., (2017) and Scholler *et al*., (2019) but using a new experimental approach, called Dynamic Full Field Optical Transmission Tomography (D-FFOTT) that is described in detail in Thouvenin *et al*., 2021. In brief, the sample is illuminated in transmission by the incoherent light emitted by a LED; this beam is partially scattered by the sample structure and partially transmitted. Both beams propagate along the same path and interfere with a phase shift (known as the Gouy phase shift) that depends on the relative position of the scatterer and of the focus of the objective. The subcellular organelle movements induce a time variation of the signal detected on each pixel of the camera. We use a Photonfocus A1024B camera working at 140 Hz with a CMOS–image– sensor with a 0.2 Me^-^ full well capacity that insures shot noise limited measurements with a good signal-to-noise ration.

To represent the evolution of signal over time we displayed a number of characteristic images extracted from a movie of typically 100 to 200 images: 1) The transmission image is analysed in order to check the stability of the cells and the lack of bleaching. 2) The average of standard deviation by group of one tenth of the movie in order to buffer any transient event. Steps 1 and 2 are run with Fiji (Schindelin et al., 2012). 3) The average cumulative sum by group of one tenth of the movie (Scholler, 2019) is compared to standard deviation of each pixel (voxel). Unlike a pure Brownian motion, a biased Brownian motion is hyper-diffusive. A way to distinguish the two types of motion is to compare the results of standard deviation to the cumulative sum. Indeed summing Brownian steps will give a trajectory that stays close to zero whereas if there is a small bias added to the Brownian random steps (**figure S1**), it will be summed for every sample and could be revealed by the cumulative sum; a more detailed description of this approach can be found in Scholler (2019). We used the cumsum function of Matlab which, when a small bias is present, is more adapted to reveal anomalous Brownian motion and, in addition increase the signal-to-noise ratio of the DCI. The normalization was made using the shot noise normal distribution in the background. All experiments have been repeated 3 times.

### Fluorescence microscopy

We used several GFP (green fluorescent protein) constructions to characterize the dynamic location of several *Phaeodactylum tricornutum* organelles. C-terminal GFP constructs were assembled by Gibson assembly into pPhat vectors, and introduced into wild-type *P. tricornutumm* by biolistic transformation, followed by selection on zeocin-supplemented ESAW-agar plates, as previously described (Falciatore et al. 1999). We used, GFP with no targeting signal, here the signal accumulates in the cytosol. GFP fused to BiP protein (Binding immunoglobulin protein) signal peptide here GFP is localized in the ER (Apt et al 2002). GFP fused to the N-terminal extension comprising a signal peptide and a transit peptide of Hsp70 of *Phaeodactylum tricornut*um (Gould et al 2006) is expressed in the PPC (periplastid compartment). Full construct vector sequences are provided in **Table S1**. For all GFP experiments transgenic diatoms were analysed by standard inverted epifluorescence microscope (Nikon Ti-E) equipped with an oil immersion objective (90X, 1.4 NA) and an EMCCD camera (Andor Ixon Ultra X-10381). GFP was excited at 488 nm and emitted fluorescence detection was from 500 to 550 nm. The sensitivity of the camera used for bright field recording (10 times lower full well capacity than the Photon Focus CMOS camera) did not allow merging of the two types of signal (fluorescence and interferometry) using the same field of view.

We also used multimodal imaging combining interferometry and fluorescence in a single optical setup (described in Thouvenin et al. 2017) to compare the fluorescent labelling with the distribution of DCI signals within lipid bodies. Diatoms were incubated with BODYPY 505/515 (dissolved in DMSO; 0.1 µg mL^-1^ final concentration) for 30minutes and observed by DCI and fluorescence. In this system we used a LED source centered at 470 nm (M470L4, Thorlabs, Newton, NJ, USA) for the excitation and filtered with a bandpass filter centered on 475 nm with a bandwidth of 50 nm (Semrock FF02-475 / 50-25).The interferometry signal was recorded on a custom camera (Quartz 2A750, Adimec). The emitted fluorescence was filtered with another bandpass filter centered on 540 nm (Semrock FF01-540 / 50-25) then imaged using a camera sCMOS (Hamamatsu Photonics). Excitation and fluorescence wavelengths were separated by a dichroic mirror at 506 nm (Semrock FF506-Di03-25).

Experiment has been repeated two times with similar results.

## RESULTS

### DCI reveals the dynamics of diatoms organelles

Diatoms grown in supplemented artificial seawater were embedded into agar in the same medium and immediately observed with the DCI set up (or D-FFOTT Thouvenin *et al*., 2021). The preparation was illuminated for a few seconds (pulsed light, 1ms, LED blue 455nm) at a few tens of Hz and a film recorded synchronously at the same frequency. We used pulsed illumination to freeze movements within cells and to avoid light saturation of photosynthesis. When observing the film of the transmission, no overall movement was detected and the diatoms looked still (**movie S1**). When we computed the standard deviation of each voxel for all the frames of the film we observed two sites of movement within the cells: vesicles or droplets and the chloroplast membrane. The following figures display the cells metabolic activity (Apelian et al., 2016) the colour map code (ImageJ, 16 colours) is such that red corresponds to the highest activity and dark blue to the lowest activity that is linked to the speed of the scatterers movements. In the vicinity of the chloroplast, vesicles or lipid droplets of different sizes, appeared green, yellow and red with the most dynamic regions located at edges of the droplet (**figure 1A**, red arrow). The other highly dynamic location corresponded to the chloroplast membranes and often looked punctuated (**figure 1A**, white arrows). Interestingly, these results were observed independently of the three morphotypes that were present in the cultures (fusiform, triradiate, or oval) (De Martino *et al*., 2007). *P. tricornutum* cells were also illuminated with a green LED (505nm) or a deep red LED (735nm, a spectral region where no *P. tricornutum* pigments absorb and no photosystems are excited). We observed with the green LED dynamic region surrounding the chloroplast and strong signal at the edges of the vesicles very similar to what we detected with the 455nm LED (**figures 2, S2A**). Histogram profiles of the standard deviation of signal intensity of each pixel were very similar between blue and 505nm LEDs with a shoulder corresponding to the strongest signals (**figure S2B)**). When we used the 735nm LED the signal was significantly reduced about 3 times (**figures 2, S2A, S2B**). We used two inhibitors at concentrations inhibiting photosystem II: 3-(3,4-dichlorophenyl)-1,1-dimethylurea (DCMU; 40 µM) or hydroxylamine (HA; 2mM) and measured the number of pixels with dynamic behaviour. We observed a significant reduction of about 3 times in presence of the inhibitors suggesting that the dynamic structures we described depend on chloroplast metabolic activity (**figure 2)**. To further investigate this observation, we compared image acquisition between continuously or pulsed LED lighting (**figure 1**). No obvious dynamic structures within the cell were highlighted when illumination was continuous unlike with pulsed light although no difference in transmission was observed meaning that no photobleaching was involved (**figures 1A and 1B**). This suggested that the high continuous light photon budget prevents chloroplast function unlike the pulsed mode level of irradiation (average power 30 times lower, **figures 1B)** and is in good agreement with the photon budget given by Prins *et al*., (2020). Our observations suggest that during the continuous exposure to light, chloroplasts are likely saturated with photons, which results in the low metabolic activity recorded (Bailleul *et al*., 2011). We reasoned that the elimination of a harmful excess energy could be dissipated through heat, however using a micro-thermocouple immersed in the sample cuvette (volume 5 µl) we did not detect an overall measurable increase in temperature (<0.1°C) under the same illumination level.

**Figure 1:**
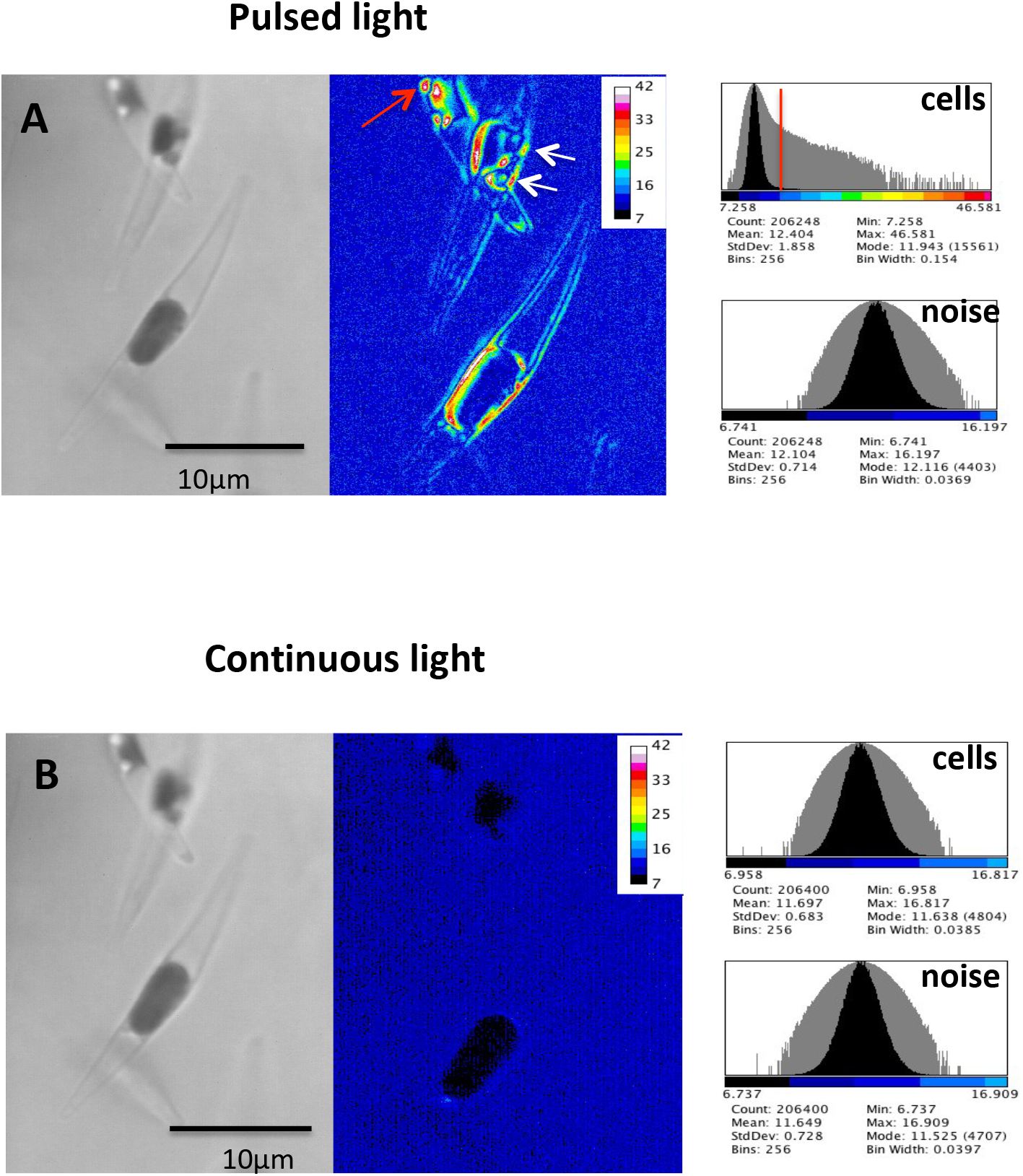
Effect of continuous or pulsed lighting on intracellular movements in *Phaeodactylum tricornutum*. The cells were successively illuminated with the LED_455_, with either continuous or pulsed light and a film of few seconds recorded. The same field in the successive acquisition was selected; left image: transmission and right image: standard deviation (scale on right). A: pulsed lighting; B: continuous lighting. Histograms of the standard deviation images using image J (Fiji) were recorded and the number of pixels of each image were computed. A: histogram of standard deviation with pulsed lighting (cells: 5638 pixels with dynamic behaviour) and histogram with pulsed lighting (noise). B: histogram of standard deviation with continuous lighting (cells: 13 pixels with dynamic behaviour) and histogram with continuous lighting (noise). Scale bar value was deduced from the format of the image (60×60 µm).

**Figure 2:**
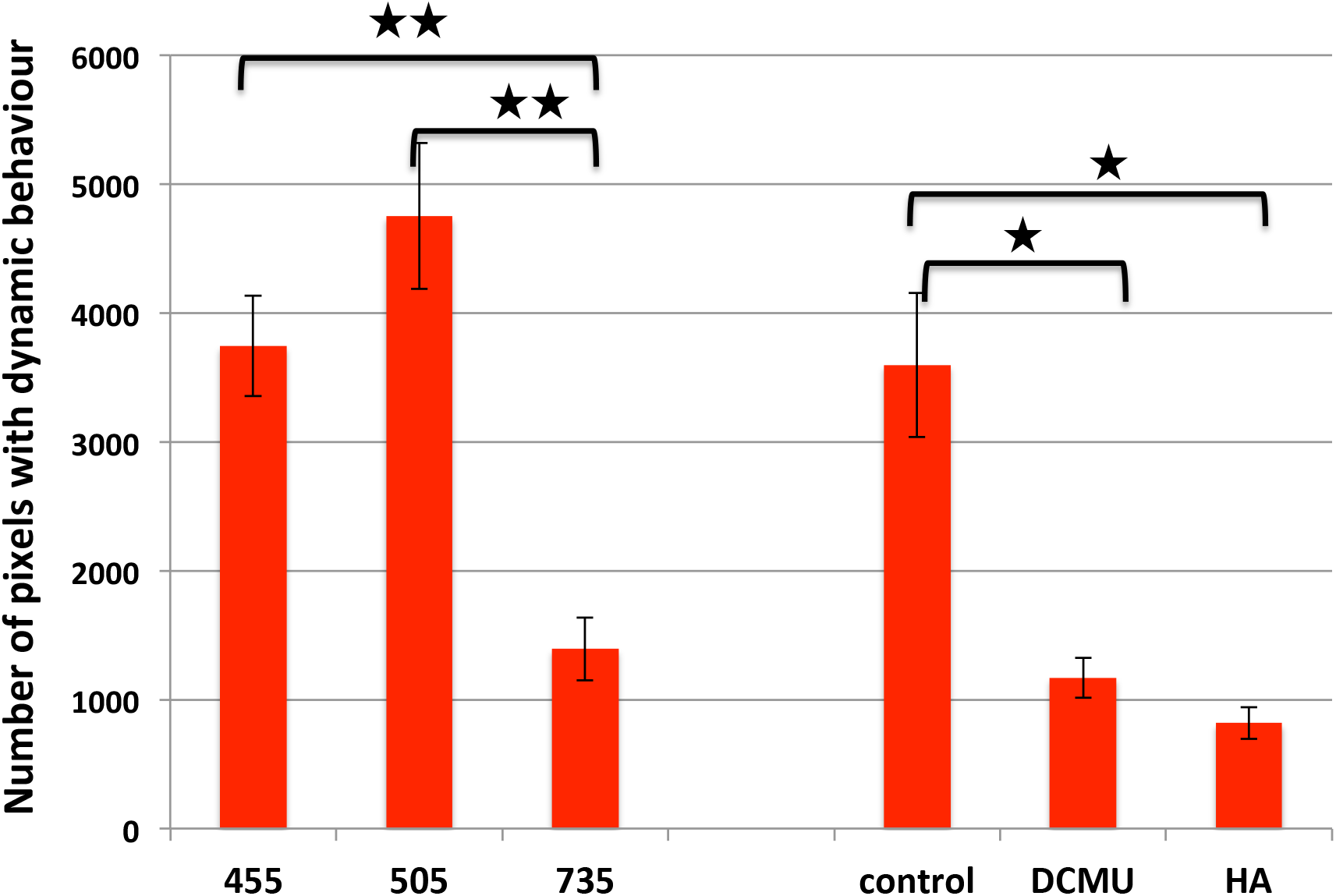
Histogramm of pixels with dynamic behaviour: effects of wavelength illumination and photosystem II inhibitors. The average values plus and minus standard deviation are presented. The different wavelengths tested are: 455nm (blue), 505nm (green) and 735nm (deep red). Two photosystem II inhibitors were used: DCMU (3,4-dichlorophenyl)-1,1-dimethyl-urea; 40µM) and HA (hydroxylamine;2mM). 20 to 30 cells were analysed. A Wilcoxon test was realized the signal observed between 455nm (blue) and 735nm (deep red) and between 505nm (green) and 735nm (deep red) gave a pvalue of 0.0079 at a significance level of 0.05) The same test was realized with the photosystem II inhibitors: The signal observed between cntrol and DCMU or HA gave a pvalue of 0.03 at a significance level of 0.05

### Fluorescence studies

To better characterize the cell compartments of the dynamic structures we described, C-terminal GFP fusions of proteins targeted to various compartments within *P. tricornutum* cells and observed with a fluorescent microscope (same samples analysed with DCI). GFP with no targeting signal led to its localization in the cytosol (**figure S3, CYT**). GFP fused to a bipartite targeting signal (signal sequence and transit peptide) of HSP70 enabling this protein to cross the two chloroplastic membranes, did not label the vesicles suggesting that they were not associated to periplastidal compartment (PPC) (**figure S3, PPC**). The signal sequence of BIP protein fused with GFP resulting in fluorescence in the endoplasmic reticulum (ER) labelled the dynamic vesicles (**figure S3, ER**). These observations suggested that vesicles possibly lipid droplets (see below) originate from the endoplasmic reticulum as already suggested by Jaussaud *et al*. (2020).

We took advantage of a setup which combined interferometry and fluorescence (Thouvenin et al. 2017) to compare the location of dynamic signals with those identifying lipid bodies. For this purpose we labelled lipid bodies with BODIPY505/515 and as shown in **figure 3** we observed co localization of dynamic signals and lipid labelling. In addition we showed in other cells that dynamic signals lighted up more structures than the specific signal observed by fluorescence (**figure S4**)

**Figure 3:**
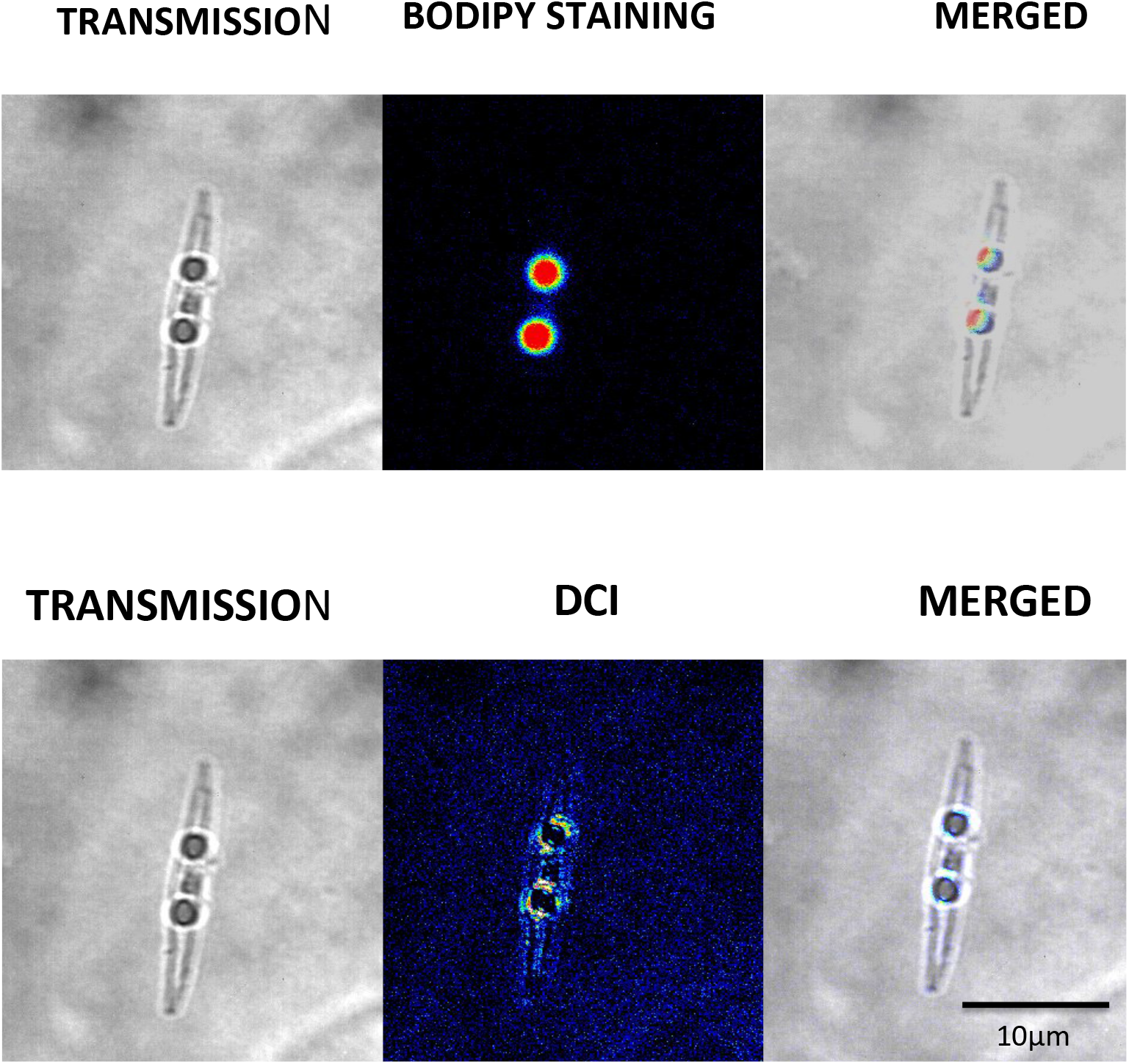
Co-localisation of dynamic droplets from *Phaeodactylum tricornutum* and BODIPY© labelled lipid droplets. Left panel: transmission microscopy, central panel: fluorescence (grey scale camera, green coloration), right panel merge of the two previous images. Left panel: transmission microscopy, central panel: Standard deviation (grey scale camera, 16 colors coloration), right panel merge of the two previous images. Scale bar value was deduced from the format of the image (90×90 µm).

### Dynamic of organelles in diatoms grown in iron-limited conditions

Kazamia *et al*. (2018) demonstrated that iron uptake in *Phaeodactylum* when grown in iron-limited conditions, involved an endocytosis step and a delivery step in the vicinity of the chloroplast. This study suggested dynamic events in *Phaeodactylum* cells when iron is limited. We thus visualized the effect of iron limitation on *P. tricornutum* intracellular dynamics after one week and two weeks of culture. After one week of culture in iron-depleted medium, a slight decrease in growth was observed suggesting a growth slow-down and after two weeks a ten-fold growth reduction was observed (**Table 1**). Samples were analysed using DCI after one (T1) or two (T2) transfers of culture with iron or iron-depleted media. In the presence of iron in the culture medium, we did not observe much difference between the two subcultures with dynamic pixels localized mostly in lipid droplets. We computed the number of pixels after removing the background noise of the same stack and found no differences between the two measurements (**Table 1, figure S4**). After one week of culture in iron-limited medium we observed a slight reduction (about 10%) in the number of pixels with dynamic behaviour. After two weeks of culture in iron-limited medium the number of pixels with dynamic behaviour were 75% reduced (**Table 1, figure S4)**. The pixels with dynamic behaviour were localized in the lipid droplets however on average there was no significant difference in the droplets diameters between the two conditions, with and without iron (1.1 ± 0.5 µm and 0.9 ± 0.2 µm respectively). All these results suggest that in the iron-depleted medium the accumulation of lipids in droplets is arrested after prolonged starvation possibly due to the shutdown of photosynthetic carbon assimilation and/or lipid synthesis.

**Table 1:** Effect of iron limitation after one (T1) and two (T2) passages on dynamic movements within *Phaeodactylum tricornutum* (*Pt*)

|  | **number of cells**<br>**10<sup>6</sup>\*mL<sup>-1</sup> ± SD** | **Average number of**<br>**pixels with dynamic**<br>**behaviour ± SD** | **Number of**<br>**analysed cells** |
| --- | --- | --- | --- |
| **+ Fe T1** | **106 ± 0.8** | **1312 ± 280** | **39** |
| **+ Fe T2** | **127 ± 2** | **1307 ± 343** | **36** |
| **- Fe T1** | **89 ± 4** | **1166 ± 311<sup>a</sup>** | **40** |
| **- Fe T2** | **10 ± 0.8** | **279 ± 131<sup>b</sup>** | **46** |

<sup>a</sup> not significantly different from +Fe T1; p-value: 0.56 (significance level of 0.05)
<sup>b</sup> significantly different from +Fe T2; p-value: 1.2 10<sup>-8</sup> (significance level of 0.05)

### Dynamics of lipid bodies in diatoms subjected to phosphate deprivation

It has been previously reported that after 8 days of culture in full nutrient-supplied media small lipid droplets start forming in *P. tricornutum* cells, suggesting the occurrence of a developing nutrient stress (Cruz de Carvalho et al., 2016). After 8 days in full culture media we observed on average, two lipid droplets per *P. tricornutum* cell and their diameters were computed (**Table 2, figures 4A, S5**). The analysis of the standard deviation of signal intensity of each pixel showed that 30 to 50% of the droplets exhibited a dynamic behaviour (**Table 2, figures 4A, S5)**. In the case of *P. tricornutum* grown for 8 days in phosphate-depleted medium there was a net reduction in cell growth (2 to 3 times) (Cruz de Carvalho et al., 2016) and a net increase in the number of droplets (5 to 10), which were significantly larger than when grown in Pi-replete medium (**Table 2**). Furthermore, the majority of the lipid droplets presented a dynamic behaviour (**Table 2, figures 4B, S5)**. Generally the droplets were in close association with the chloroplast but sometimes they were found free in the cytoplasm and were able to fuse giving rise to very large structures (red arrow, **figure S5**). Histogram profiles showed a net increase (about 5 fold) in the number of pixels above the background noise between the conditions with and without phosphate (**Table 3, figures 4, S5)**.

**Table 2:** Diameter and dynamic behaviour of droplets in full or phosphate-depleted medium.

|  | **Average droplet diameter ($\mu\text{m} \pm \text{SD}$)**<br>**(number of analysed droplets)** | **Number of droplets with dynamic behaviour**<br>**(percentage)** |
| --- | --- | --- |
| **+ Pi: exp1** | **0.56±0.22 (42)** | **22 (52%)** |
| **- Pi: exp1** | **0.97±0.33 (58)** | **55 (94%)** |
| **+ Pi: exp2** | **0.43±0.11 (18)** | **6 (33%)** |
| **- Pi: exp2** | **0.96±0.32 (26)** | **21 (80%)** |
| **+ Pi: exp3** | **0.51±0.21 (26)** | **12 (46%)** |
| **- Pi: exp3** | **0.88±0.22 (86)** | **61 (71%)** |

**Table 3:** Comparison of movements of the *Phaeodactylum* scatterers computing their standard deviation or cumulative sum in media with or without phosphate.

|  | **Number<sup>a</sup> of**<br>**dynamic pixels**<br>**±SD (method:**<br>**standard**<br>**deviation)** | **Number of**<br>**dynamic pixels**<br>**±SD (method:**<br>**Cumulative**<br>**sum)** | **Ratio**<br>**Cumulative sum/**<br>**Standard deviation** |
| --- | --- | --- | --- |
| **+Pi** | **586±167** | **1174±196** | **2** |
| **-Pi** | **2844±155** | **3311±236** | **1.16** |

<sup>a</sup> Three stacks have been analysed in each conditions

**Figure 4:**
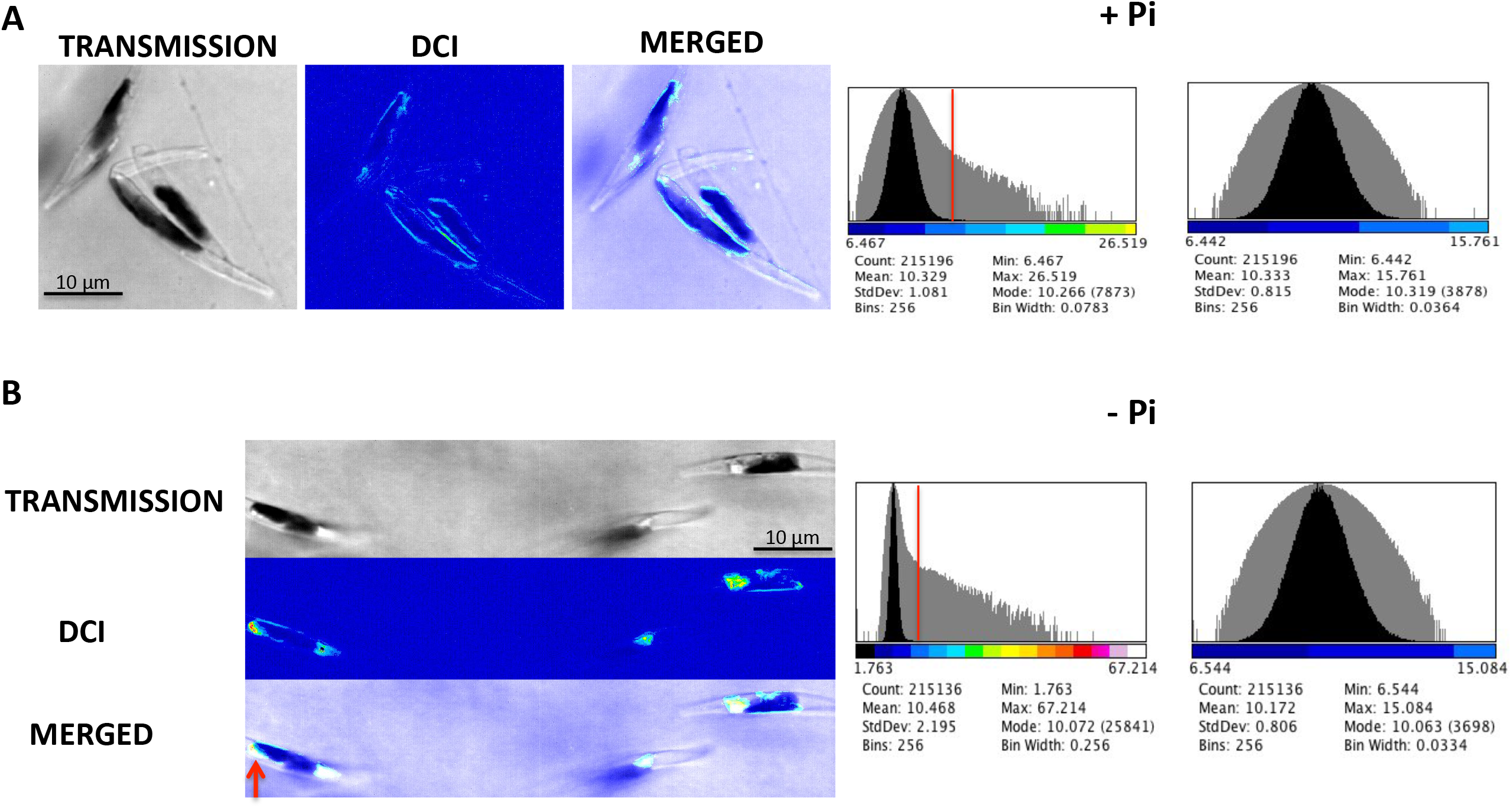
Increase in lipid droplets size and numbers in *P. tricornutum* cells grown for 8 days in phosphate-depleted medium. From left to right panel (or from top to bottom) are shown, the transmission, the standard deviation of each pixel and the merged image (artificial blue colour). Histograms of the standard deviation images using image J (Fiji) were recorded and the number of pixels above noise level were computed (indicated by a red bar on the histogram). Histograms of the same image size without cells, were recorded to determine noise levels. Scale bar value was deduced from the format of the image (60×60 µm.)

To get more insights into the dynamics of lipid droplets we analysed the time series of each pixel in order to get a map of the transmission, the standard deviation, the cumulative sum (Matalb: cumsum function) and the average frequency. Despite a noticeable difference in signal level (standard deviation and cumulative sum, see below) between the cells grown with Pi and without Pi, we did not observe any significant differences with regard to the central frequencies, which were around 15 +/-5 Hz. We observed a 3-5 times increase in the signal amplitude (standard deviation and cumulative sum respectively) in *P. tricornutum* grown without phosphate (**Table 3**). The analysis of the signal value also pointed out also some differences of the signal distribution between standard deviation and cumulative sum computations. The computation of value variation of a pixel relied on the cumulative sum of the signals highlighted a non-Brownian displacement within the lipid droplets (**Table 3**). Following the method developed in detail in Scholler (2019) we found that the normalized cumulative sum is about two times that of the normalized standard deviation in culture with phosphate while it is slightly superior (1.2 times) in culture without phosphate (**Table 3)**. This result suggests that the random movement of scatterers within the droplets is likely to be associated to a drift that makes it hyper-diffusive, rather than being a pure Brownian motion and could correspond to the flux of lipids and proteins filling lipid bodies (Olzmann and Carvalho 2019; Nettebrock and Bohnert 2020).

## DISCUSSION

Here, we present a non-invasive and non-destructive method, which allows the detection and characterisation of internal movements within photosynthetic cells. In particular, through image analysis we show evidence that these dynamic processes are correlated with the functioning of the chloroplasts. This detection was possible by relying on the scattering properties of the cell organelles and of the high frame rate of our camera recordings. We were able to identify and quantify movements within lipid droplets, specific organelles, which accumulate in nutrient-stressed diatoms. In cells deprived of phosphate in addition to the random movement of little scatterers, which may correspond to the movement of membrane proteins from ER (Jacquier et al., 2011) or storage of misfolded proteins in lipid droplets as suggested by Lupette *et al*. (2019), we described a drift movement of scatterers, which might correspond to molecular fluxes of proteins and/or lipids. The distribution of dynamic pixels often observed in an eggcup shape (**figure S6**) could be in agreement with ER surrounding lipid droplets (Jacquier *et al*., 2011) and more precisely in the specific case of diatoms the most outer membrane of the chloroplast (Flori *et al*., 2016, Leyland *et al*., 2020, Jaussaud *et al*., 2020).

The DCI method can be extended to any microalgae considering that transformation is not available for many of them. We undertook a similar study with *Fragilariopsis cylindrus* grown under light or in dark and we observed many dynamic structures within this diatom (not shown). The method is very sensitive and allows the study of slight changes due to environmental fluctuations. In addition, most of image analyses have been run with Fiji (ImageJ), a public domain application (Schindelin et al., 2012).

Cytoplasmic streaming generated by myosin in the alga *Chara* or in plants has long been described (for review, Tominaga & iIto, 2013). Indeed the speed of cytoplasmic streaming generated by *Chara* myosin observed by optical trap nanometry is in the range of 50 µm/s (Tominaga & iIto, 2013). With DCI we observed life imaging of low speed movements (about 0.1 µm/s) of undetectable scatterers within *P. tricornutum* organelles and corresponding to their metabolic activity. This method could be extended to other dynamic structures such as liquid droplets present in other organelles or in nuclei.

## ACKNOWLEGEMENTS

We wish to thank our colleagues: Ignacio Izzedine (Institut Langevin) for the use of the fluorescence microscope, Olivier Thouvenin (Institut Langevin) for his help in the dynamic signal analysis, and Benjamin Bailleuil for providing photosystem II inhibitors. HB and CB acknowledge financial support from the DIM ELICIT (INFLUERE IN VISCERA). FC acknowledges Q-Life for the PhD fellowship (Q-life ANR-17-CONV-0005), HCC acknowledges funding from the Agence National pour la Recherche (ANR DiaLincs 19-CE43-0011-01), RGD acknowledges a CNRS Momentum Fellowship. CB acknowledges funding from the ERC (Diatomic project) and the ANR (Browncut).

## FUNDING

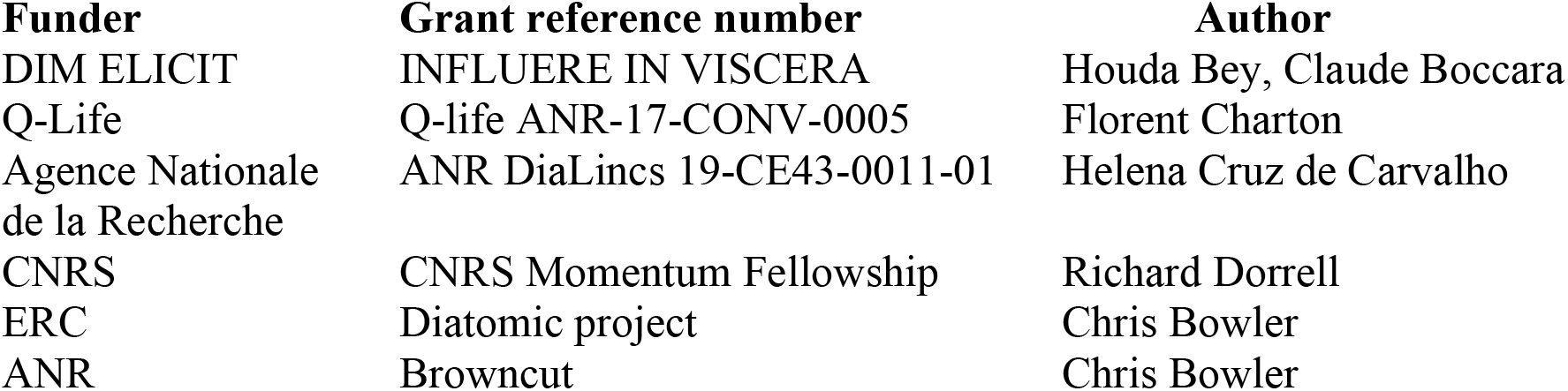

## SUPPORTING INFORMATION

### SUPPLEMENTARY TABLE

**Table S1: list of primers for GFP constructs**

### SUPPLEMENTARY MOVIE

**Movie S1: Transmission film of diatoms illuminated at 455nm**

### SUPPLEMENTARY FIGURES

**Figure S1: Simulation of a pure Brownian motion (blue) and of a biased movement (red) where a drift is added to the Brownian motion**.

The y-axis of the graph shows. the dynamic movement of a particle) measured over time (x-axis). For this specific example the standard deviation was found to be 1.16 (Brownian) and 1.37 (Brownian biased) whereas the cumsum (cumulative sum) was found to be 12.30 (Brownian) and 180.64 (Brownian biased).), i.e. demonstrating a the very large enhancement of the signal when a drift is present.

**Figure S2: Effect of wavelength illumination on intracellular movements in *Phaeodactylum tricornutum***

Cells were successively illuminated with pulsed light from LED_735_ (red), LED_505_ (green) and LED_455_ (blue) and a film of few seconds recorded. The same field of view in the successive acquisition was selected; upper image: transmission; lower image standard deviation (scale on bottom left) **(figure S2A)**. Histograms of the standard deviation images using image J (Fiji) were recorded and the number of pixels above noise level were computed (indicated by a red bar on the histogram) in **figure S2B**. Histograms of the same image for each wavelength, but without cells, were recorded to determine noise levels. Scale bar value was deduced from the format of the image (60×60 µm)

**Figure S3: Co-localisation of dynamic lipid droplets from *Phaeodactylum tricornutum* and labelled endoplasmic reticulum**

Left panel: light microscopy, central panel: GFP fluorescence (grey scale camera, green coloration), right panel merge of the two upper images. Cytosolic (CYT) expressed GFP accumulates in the entire cell. The signal peptide of BIP protein fused to GFP leads to endoplasmic localization of GFP (ER). The dual topogenic signal of Hsp70 fused to GFP targets GFP to periplastid compartment (PPC). Scale bar value was deduced from the format of the image (60×60 µm).

**Figure S4: Dynamic signal in Phaeodactylum is not limited to lipid bodies**

Upper panel: transmission microscopy, central panel: Standard deviation (grey scale camera, 16 colors), lower panel merge of the two previous images. Scale bar value was deduced from the format of the image (90×90 µm).

**Figure S5: Decrease in dynamic signal in *Phaeodactylum tricornutum* grown in iron depleted medium**

Representative cells grown in medium with iron (+Fe) or depleted of iron (-Fe). From left to right panel is shown the transmission, the standard deviation of each pixel and the merged image (artificial blue colour). T1 and T2 correspond to 8 days and 16 days growth in medium with iron or depleted of iron. Histograms of the standard deviation images using image J (Fiji) were recorded and the number of pixels above noise level were computed (indicated by a red bar on the histogram) in **Table 1**. Histograms of the same image size without cells, were recorded to determine noise levels Scale bar value was deduced from the format of the image (60×60 µm)

**Figure S6: Diatoms grown with and without Phosphate: Increase in lipid droplets size and numbers in medium depleted in Phosphate**

## AUTHOR CONTRIBUTIONS

M.B. performed, and analyzed the biological experiments and wrote the manuscript. C.B. built, characterized the optical setup and analyzed the movies and wrote the manuscript. HB and FC compared interferometry and fluorescence. SL and RD built the GFP fusions. FC and SD performed the culture of *P tricornutum* and studied their growth under stress. RD, HCC and CB contibute the writing of the manuscript. All authors discussed the results and commented on the manuscript.

## Notes

### Competing Interest Statement

The authors have declared no competing interest.

