## supplementary figures for "Dynamic Cell Imaging: application to the diatom *Phaeodactylum tricornutum* under environmental stresses"

### Table S1

| Vector | Localisation | Insert |
|---|---|---|
| **pPhat eGFP** | **Cytoplasm** | |
| **BiP NTD eGFP** | **CER** | ATGATATTCATGAGAATTGCCG<br>TAGCAGCACTGGCCTTGCTGGC<br>TGCTCCCTCCATTCGTGCCGAAG<br>AGGCCGGTGAAGAGGCCAAGA<br>TGGGTACCGTG |
| **Hsp70 NTD eGFP** | **PPC** | ATGGTGCATCTTCCATCCTCCTC<br>CACCCTCCTAGCCTGCGTGTCG<br>GTACTTCTCTCCGGAGCTCACCC<br>CGCAAAAGCATCATGGCTAGCT<br>CGTAGGACAGTAGAAAAGCCG<br>ACTCTGGCACGAATCCATGAGC<br>AGCGAGACTCTACGGACCGGAA<br>ATCGAGGGCGCCCTTCTTACATC<br>GTCGACAGTATCACTACGCGCA<br>TCATCCCCACGGGCTACTCACTA<br>CCCTCCGTGGTGGCGCCAGTAA<br>TGCAGACAACAAAATGGGATCC |

### movie S1

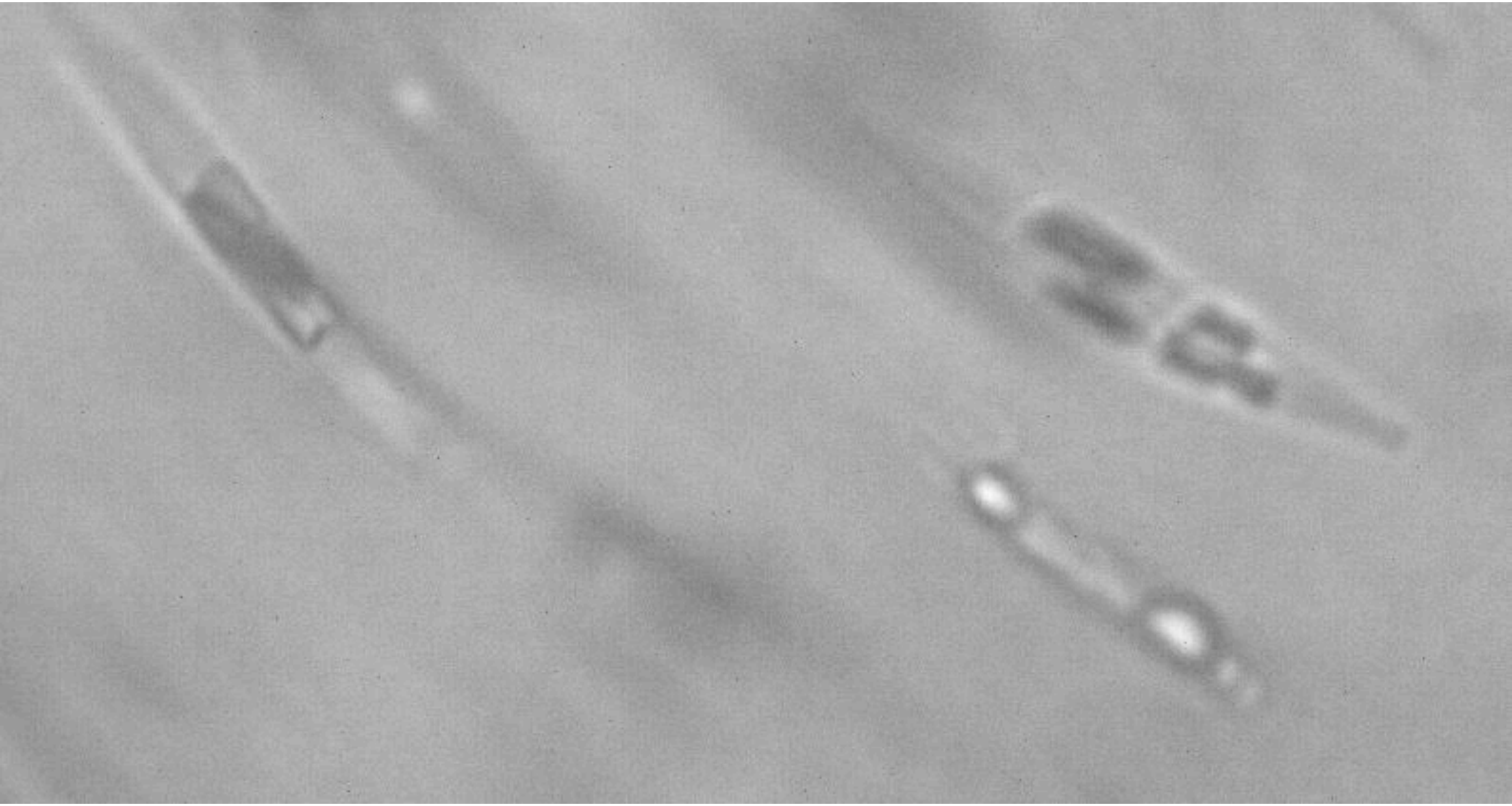

### Figure S1

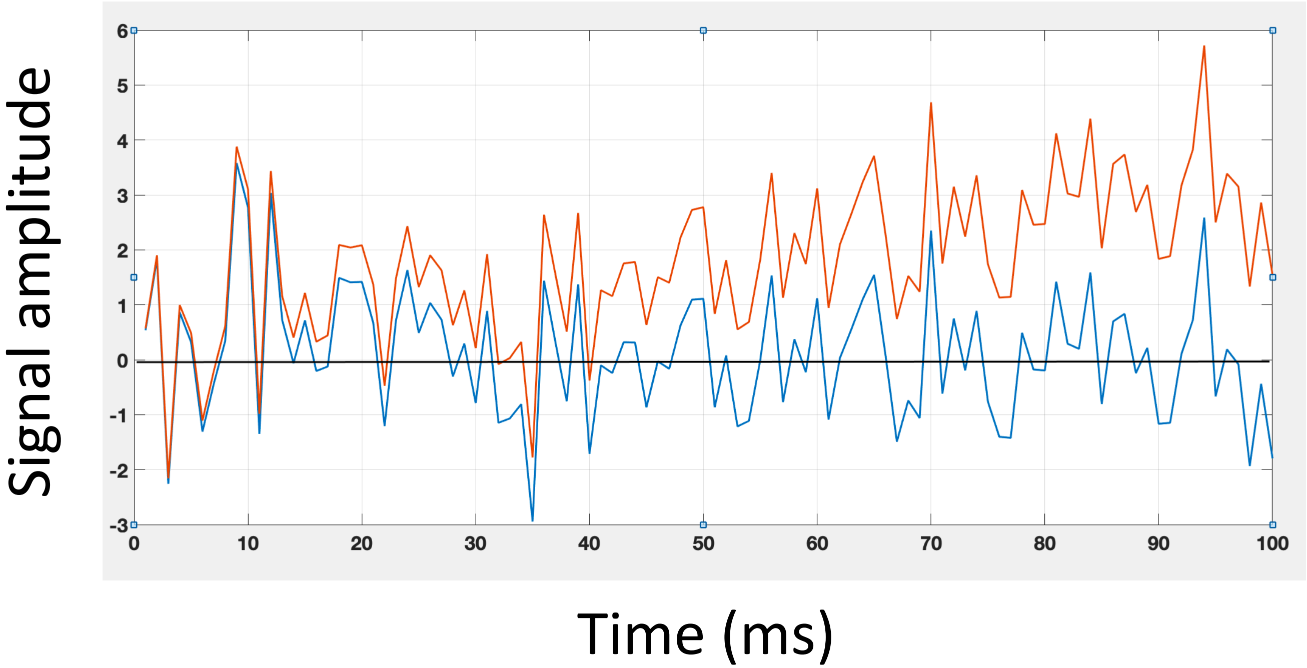

Signal amplitude
6
5
4
3
2
1
0
-1
-2
-3
0
10
20
30
40
50
60
70
80
90
100
Time (ms)

#### Figure S2A

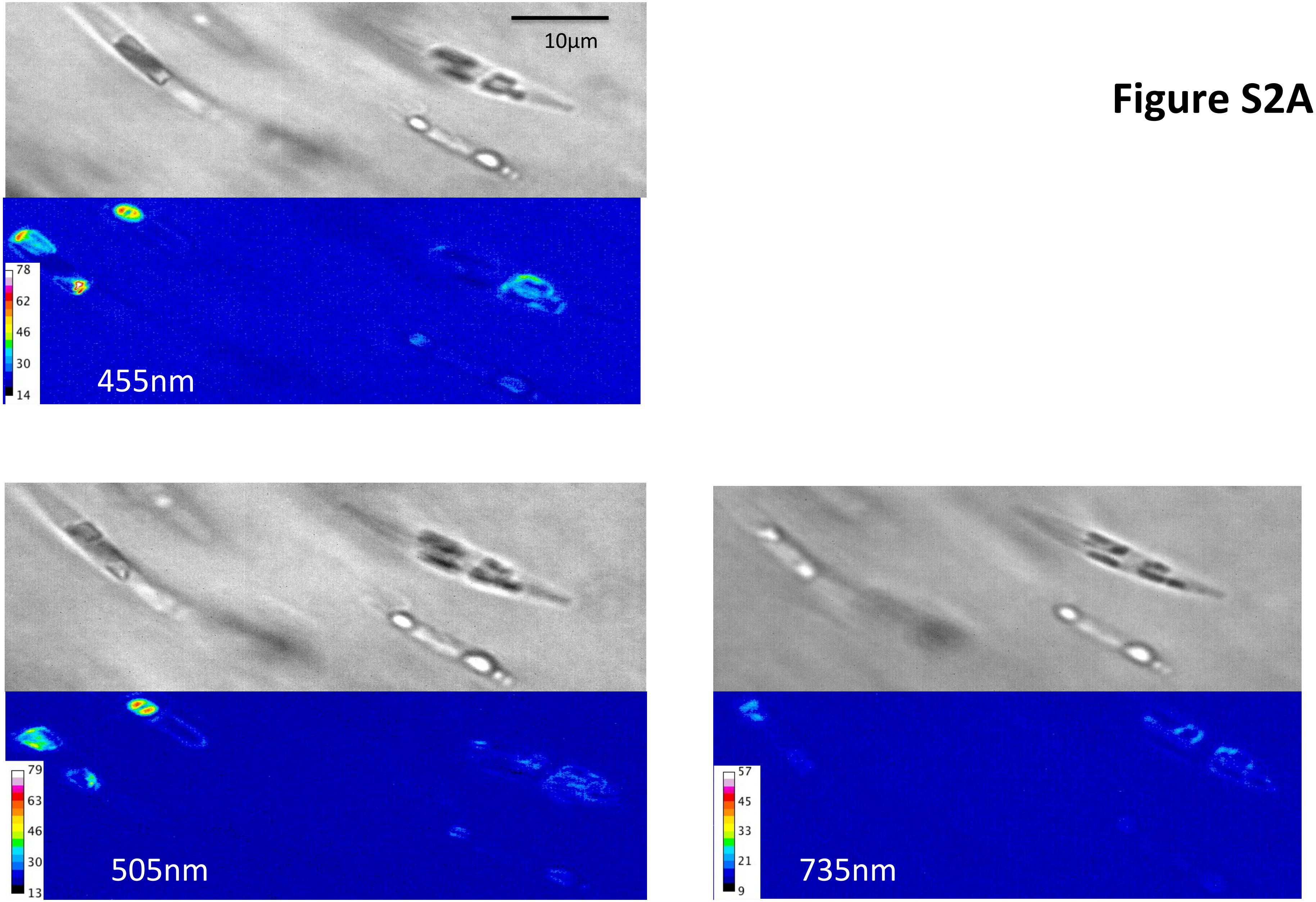

### Figure S2B

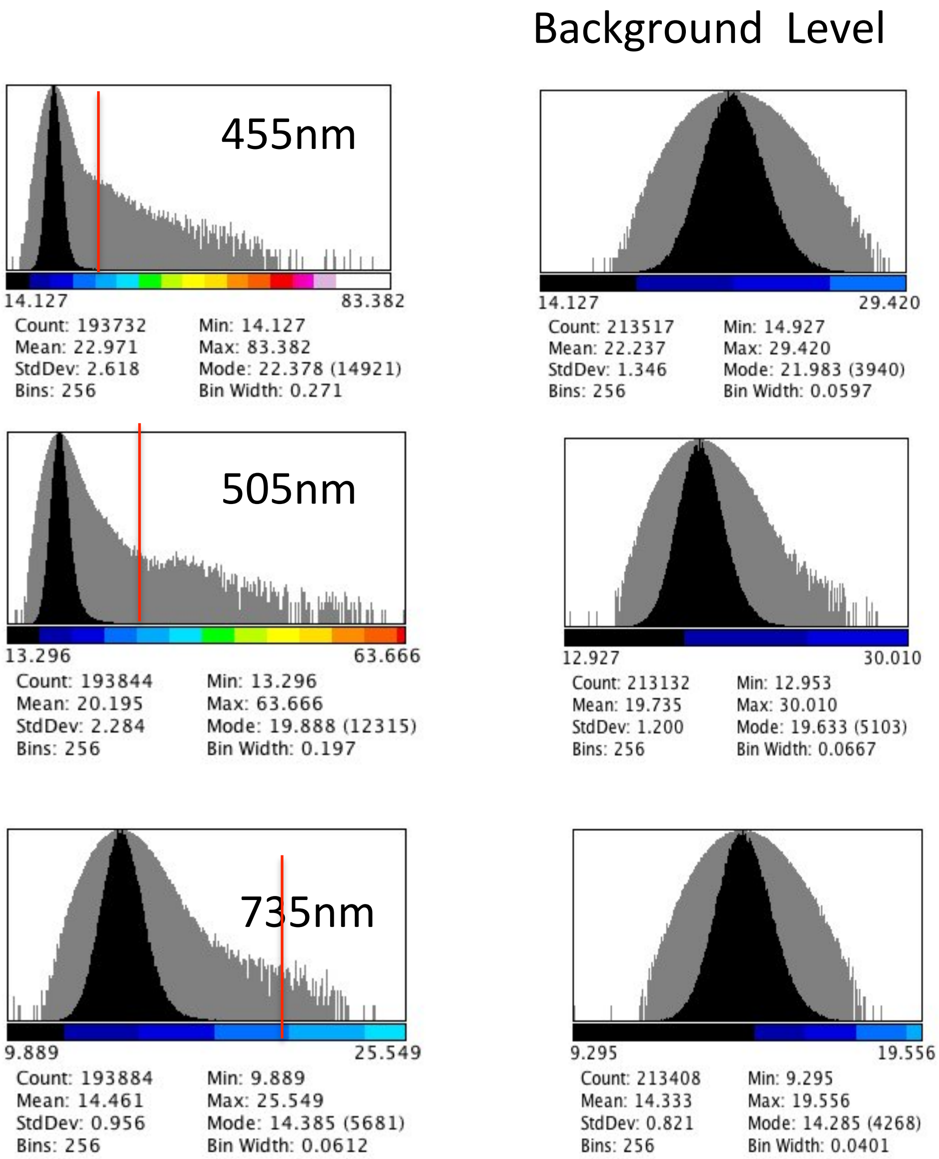

Background Level
455nm
14.127
83.382
Count: 193732
Mean: 22.971
StdDev: 2.618
Bins: 256
Min: 14.127
Max: 83.382
Mode: 22.378 (14921)
Bin Width: 0.271
14.127
29.420
Count: 213517
Mean: 22.237
StdDev: 1.346
Bins: 256
Min: 14.927
Max: 29.420
Mode: 21.983 (3940)
Bin Width: 0.0597
505nm
13.296
63.666
Count: 193844
Mean: 20.195
StdDev: 2.284
Bins: 256
Min: 13.296
Max: 63.666
Mode: 19.888 (12315)
Bin Width: 0.197
12.927
30.010
Count: 213132
Mean: 19.735
StdDev: 1.200
Bins: 256
Min: 12.953
Max: 30.010
Mode: 19.633 (5103)
Bin Width: 0.0667
735nm
9.889
25.549
Count: 193884
Mean: 14.461
StdDev: 0.956
Bins: 256
Min: 9.889
Max: 25.549
Mode: 14.385 (5681)
Bin Width: 0.0612
9.295
19.556
Count: 213408
Mean: 14.333
StdDev: 0.821
Bins: 256
Min: 9.295
Max: 19.556
Mode: 14.285 (4268)
Bin Width: 0.0401

### Figure S3

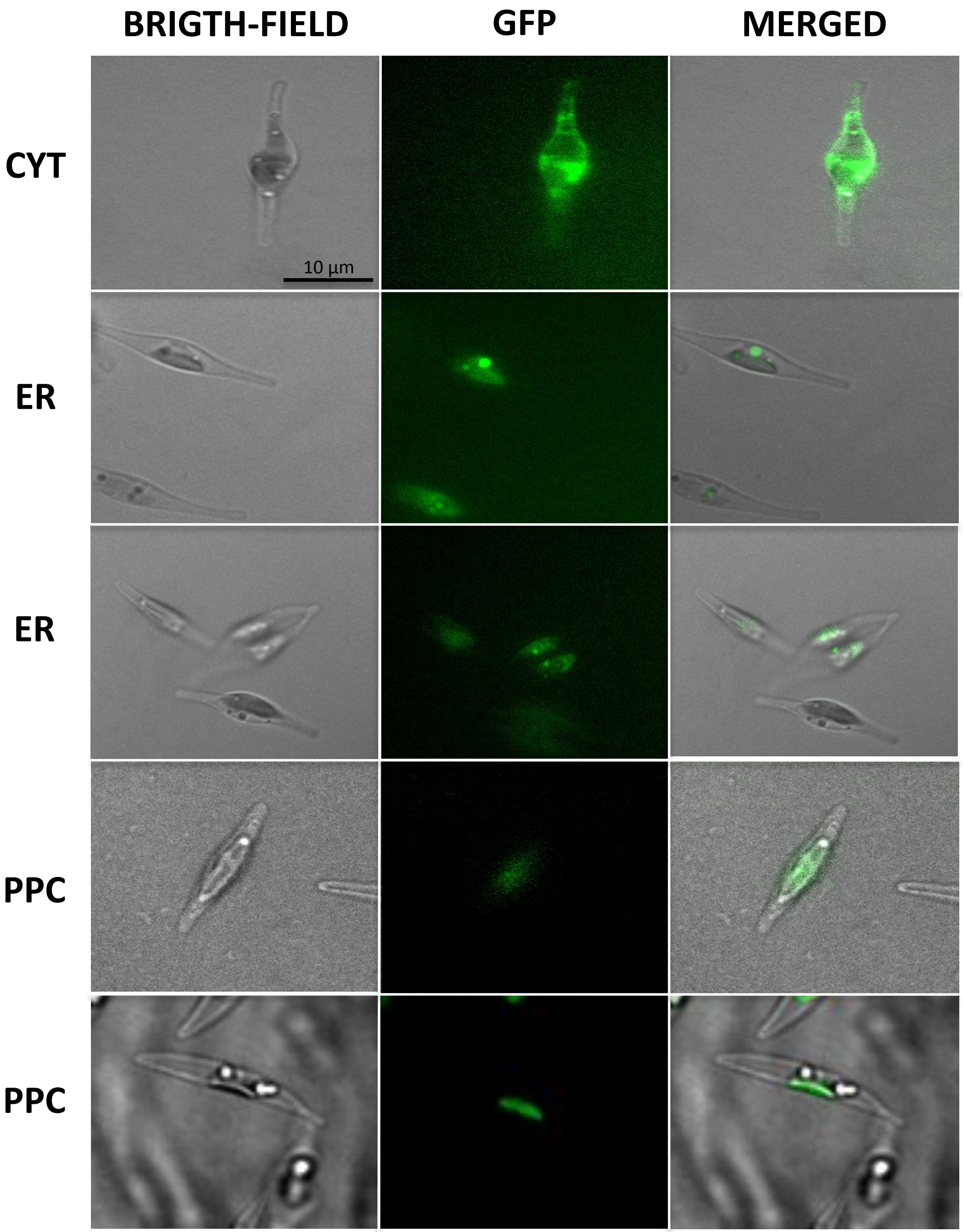

### Figure S4

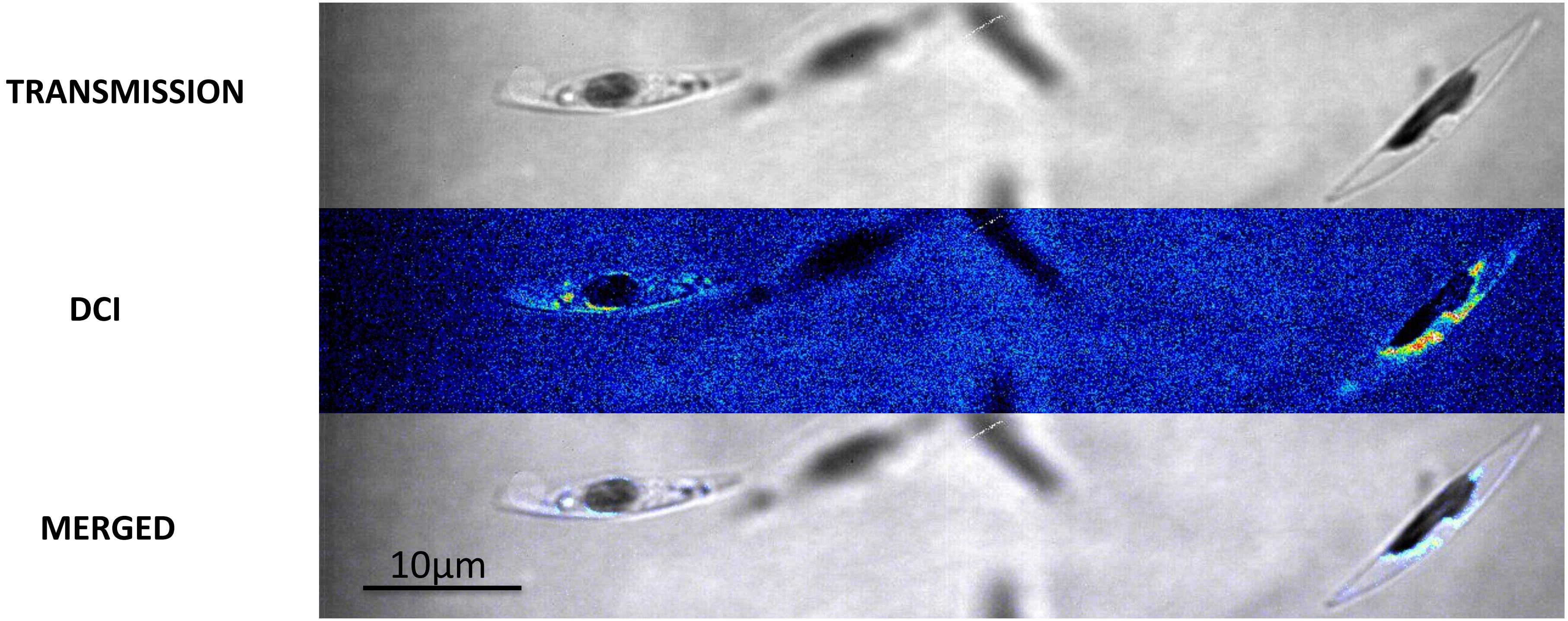

### Figure S5

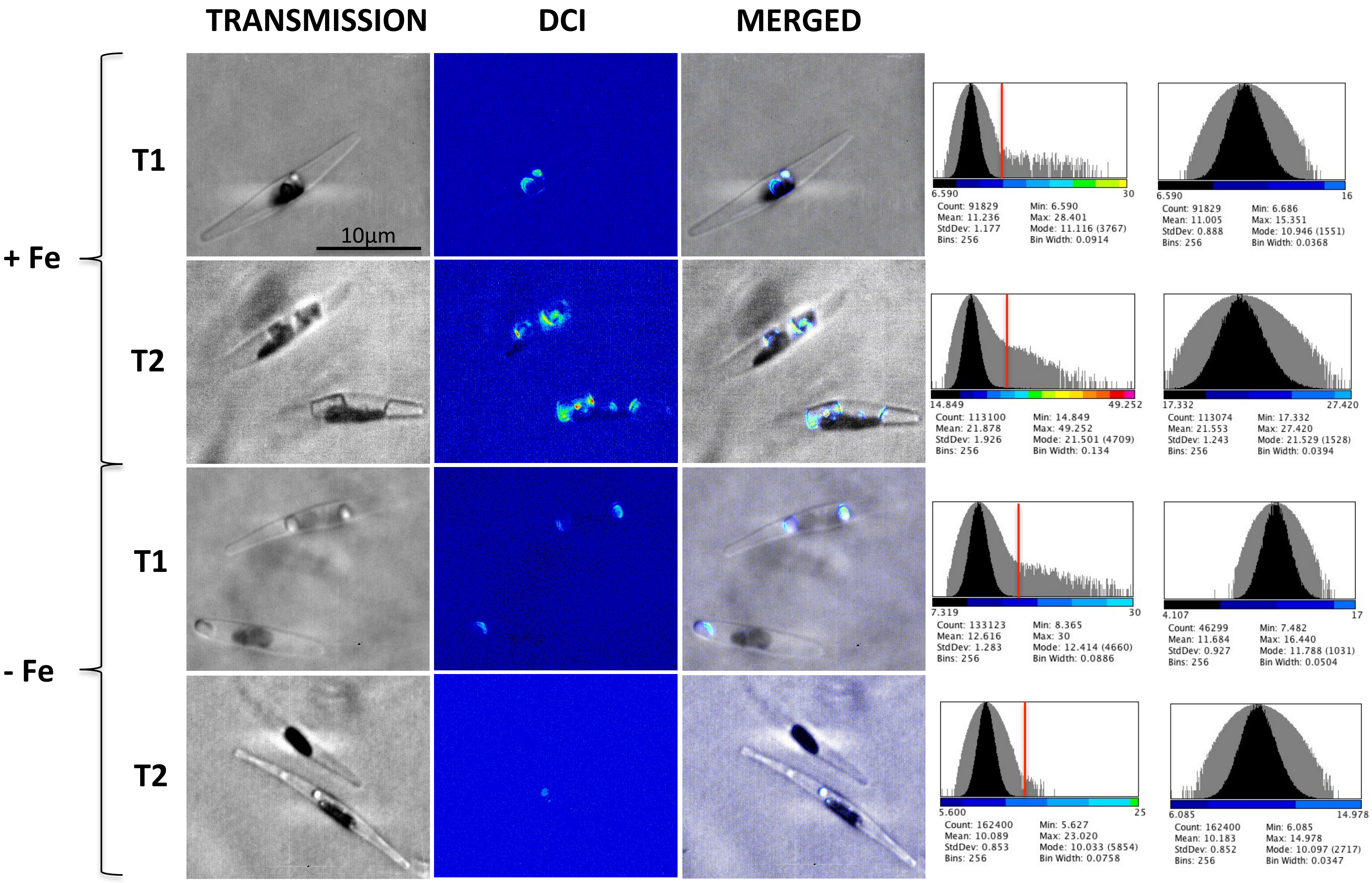

TRANSMISSION
DCI
MERGED
+ Fe
- Fe
T1
T2
T1
T2
10µm
6.590
30
Count: 91829
Mean: 11.236
StdDev: 1.177
Bins: 256
Min: 6.590
Max: 28.401
Mode: 11.116 (3767)
Bin Width: 0.0914
6.590
16
Count: 91829
Mean: 11.005
StdDev: 0.888
Bins: 256
Min: 6.686
Max: 15.351
Mode: 10.946 (1551)
Bin Width: 0.0368
14.849
49.252
Count: 113100
Mean: 21.878
StdDev: 1.926
Bins: 256
Min: 14.849
Max: 49.252
Mode: 21.501 (4709)
Bin Width: 0.134
17.332
27.420
Count: 113074
Mean: 21.553
StdDev: 1.243
Bins: 256
Min: 17.332
Max: 27.420
Mode: 21.529 (1528)
Bin Width: 0.0394
7.319
30
Count: 133123
Mean: 12.616
StdDev: 1.283
Bins: 256
Min: 8.365
Max: 30
Mode: 12.414 (4660)
Bin Width: 0.0886
4.107
17
Count: 46299
Mean: 11.684
StdDev: 0.927
Bins: 256
Min: 7.482
Max: 16.440
Mode: 11.788 (1031)
Bin Width: 0.0504
5.600
25
Count: 162400
Mean: 10.089
StdDev: 0.853
Bins: 256
Min: 5.627
Max: 23.020
Mode: 10.033 (5854)
Bin Width: 0.0758
6.085
14.978
Count: 162400
Mean: 10.183
StdDev: 0.852
Bins: 256
Min: 6.085
Max: 14.978
Mode: 10.097 (2717)
Bin Width: 0.0347

### Figure S6

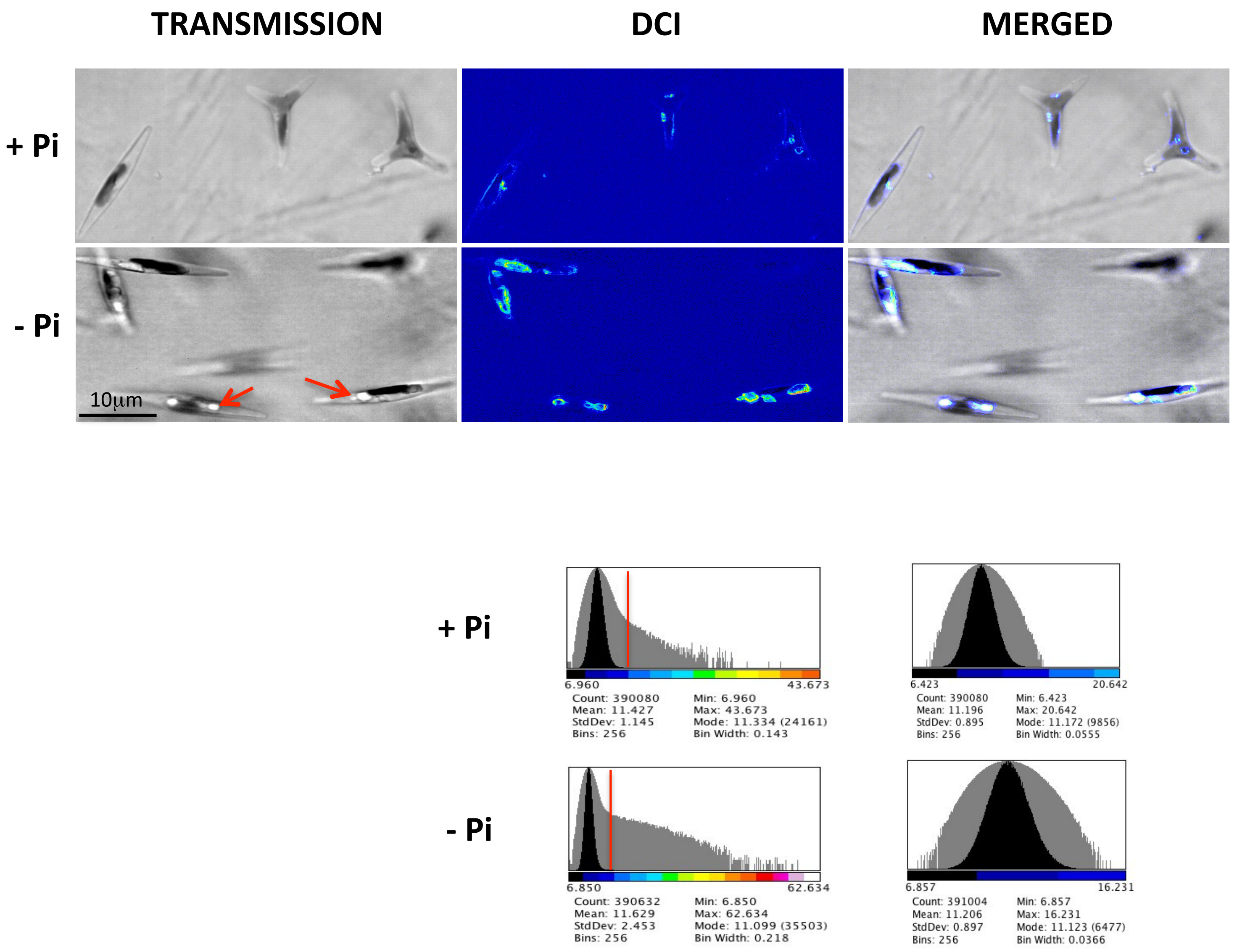

TRANSMISSION
DCI
MERGED
+ Pi
- Pi
10μm
+ Pi
6.960
43.673
Count: 390080
Mean: 11.427
StdDev: 1.145
Bins: 256
Min: 6.960
Max: 43.673
Mode: 11.334 (24161)
Bin Width: 0.143
6.423
20.642
Count: 390080
Mean: 11.196
StdDev: 0.895
Bins: 256
Min: 6.423
Max: 20.642
Mode: 11.172 (9856)
Bin Width: 0.0555
- Pi
6.850
62.634
Count: 390632
Mean: 11.629
StdDev: 2.453
Bins: 256
Min: 6.850
Max: 62.634
Mode: 11.099 (35503)
Bin Width: 0.218
6.857
16.231
Count: 391004
Mean: 11.206
StdDev: 0.897
Bins: 256
Min: 6.857
Max: 16.231
Mode: 11.123 (6477)
Bin Width: 0.0366
